# Non-Invasive Brain Stimulation to Investigate Language Production in Healthy Speakers: A Meta-Analysis

**DOI:** 10.1101/230631

**Authors:** Jana Klaus, Dennis J.L.G. Schutter

**Affiliations:** Radboud University

**Keywords:** language production, meta-analysis, picture naming, verbal fluency, TMS, tDCS

## Abstract

Non-invasive brain stimulation (NIBS) has become a common method to study the interrelations between the brain and language functioning. This meta-analysis examined the efficacy of transcranial magnetic stimulation (TMS) and direct current stimulation (tDCS) in the study of language production in healthy volunteers. Forty-five effect sizes from 30 studies which investigated the effects of NIBS on picture naming or verbal fluency in healthy participants were meta-analysed. Further sub-analyses investigated potential influences of stimulation type, control, target site, task, online vs. offline application, and current density of the target electrode. Random effects modelling showed a small, but reliable effect of NIBS on language production. Subsequent analyses indicated larger weighted mean effect sizes for TMS as compared to tDCS studies. No statistical differences for the other sub-analyses were observed. We conclude that NIBS is a useful method for neuroscientific studies on language production in healthy volunteers.

## Introduction

Transcranial magnetic (TMS) and direct current stimulation (tDCS) are non-invasive brain stimulation (NIBS) techniques that are increasingly used to investigate causal relationships between language functions and their underlying neuronal processes. The aim of this combined review and meta-analysis is to examine the efficacy and reliability of NIBS as an intervention method to study the neural correlates of language production in healthy volunteers. Prior meta-analyses on the effects of transcranial direct current stimulation (tDCS) on verbal fluency and picture naming have provided diverging results. Both Horvath, Forte, and Carter (2015) and Price, McAdams, Grossman, and Hamilton (2015) analysed performance changes in semantic production and word learning tasks, with the first finding no effect, but the latter reporting a reliable modulation of task performance. Furthermore, Westwood and Romani (2017) found no effect of tDCS on language production performance across production and reading tasks. Our present review offers an overview and meta-analysis of studies which measured changes in verbal fluency and picture-naming performance during or following the administration of tDCS or transcranial magnetic stimulation (TMS). Furthermore, by differentiating between different experimental parameters, we aim to provide a more detailed picture with respect to the usefulness of NIBS studies that investigate language production in healthy volunteers.

Picture naming (i.e., the production of a noun or verb in response to a visually presented stimulus) is the most direct way to measure language production performance. Cortical activity during this task has been located in a large left frontotemporal network stretching from the interior frontal to posterior superior temporal and inferior parietal regions (Indefrey, 2011; Indefrey & Levelt, 2004). Using TMS, which applies an ultra-short electromagnetic pulse that creates an electric current in superficial cortical nerve tissue, an engagement of the posterior superior temporal gyrus (pSTG), middle temporal gyrus (MTG), anterior temporal lobe (ATL), and inferior frontal gyrus (IFG) has been demonstrated (Acheson, Hamidi, Binder, & Postle, 2011; Mottaghy et al., 1999; Pobric, Jefferies, & Lambon Ralph, 2007, 2010, Schuhmann, Schiller, Goebel, & Sack, 2009, 2012; Shinshi et al., 2015; Sparing et al., 2001; Töpper, Mottaghy, Brügmann, Noth, & Huber, 1998; Wheat et al., 2013). Furthermore, cortical excitability can be modulated by applying a constant weak electric current between two electrodes affixed on the scalp. Although the vast majority of the electric field is shunted, a small yet significant portion of the field reaches the superficial layers of the cortex (Nitsche et al., 2008). Research on the human motor cortex has shown that anodal tDCS increases spontaneous neural firing and cortical excitability, while cathodal tDCS reduced spontaneous neural firing and lowered cortical excitability (Nitsche & Paulus, 2000; Stagg & Nitsche, 2011). Its potential to modulate underlying cortical tissue together with the facts that tDCS is not associated with serious adverse advents and allows for better (double) blinding procedures as compared to TMS has contributed to its increased use in cognitive neuroscience. Indeed, a number of studies have reported significant effects from applying anodal tDCS over the left STG and dorsolateral prefrontal cortex (DLPFC) on object and action naming (Fertonani, Brambilla, Cotelli, & Miniussi, 2014; Fertonani, Rosini, Cotelli, Rossini, & Miniussi, 2010; Sparing, Dafotakis, Meister, Thirugnanasambandam, & Fink, 2008). Interestingly, NIBS typically only affects naming latencies, but not error rates, in picture naming tasks. Next to the classic picture naming tasks, a number of studies have also investigated the effects of tDCS and TMS on naming latencies in the semantic blocking and picture-word interference paradigm. In semantic blocking tasks, naming latencies are compared between semantically homogeneous (i.e., containing words from the same semantic category) and heterogeneous blocks (i.e., semantically unrelated words). Retrieving and producing semantically related words in a row typically results in longer naming latencies compared to producing semantically unrelated words. This semantic interference (SI) effect is taken as evidence for competitive selection of target responses (e.g., Belke, Meyer, & Damian, 2005; Damian, Vigliocco, & Levelt, 2001; Kroll & Stewart, 1994) and has been localised predominantly in the left temporal cortex (de Zubicaray, Johnson, Howard, & McMahon, 2014; Indefrey, 2011). Confirming this, studies applying tDCS (Meinzer, Yetim, McMahon, & de Zubicaray, 2016; Pisoni, Papagno, & Cattaneo, 2012) or TMS (Krieger-Redwood & Jefferies, 2014) before or during semantic blocking tasks reported an involvement of pSTG, but not IFG. These studies provide first evidence that processes involving lexical selection and retrieval can be targeted using NIBS. However, it should be kept in mind that behavioral effects were numerically small (see also Westwood et al., 2017, Experiment 2, for statistical null effects of tDCS across the left IFG in a semantic blocking task).

The picture-word interference (PWI) paradigm allows for the chronometric investigation of speech production processes on the timescale of tens of milliseconds (e.g., Damian & Martin, 1999; Schriefers, Meyer, & Levelt, 1990). Participants are asked to name pictures while ignoring a visually or auditorily presented distractor word, the relatedness of which to the target word is systematically varied. Typically, a semantically related distractor (e.g., *“cow”* when the target word is *“sheep”*) increases naming latencies compared to an unrelated distractor, while a phonologically related distractor (e.g., *“sheet”*) speeds up naming latencies. Varying the onset of the distractor relative to picture presentation (stimulus-onset asynchrony, SOA) enables researchers to examine the time course of speech planning with respect to the individual representational levels involved. Recall that lexical-semantic processing has been associated with the left MTG, while phonological processing has been located in the left STG (Indefrey, 2011; Indefrey & Levelt, 2004). In line with this, Henseler, Mädebach, Kotz, & Jescheniak (2014) reported a decrease of associative facilitation (i.e., when the distractor is associatively related vs. unrelated to the target word, e.g. *“boat”* and *“port”*) under MTG as opposed to IFG and sham stimulation (anodal tDCS). Furthermore, Pisoni, Cerciello, Cattaneo, & Papagno (2017) found reduced phonological facilitation following anodal tDCS to the STG, but no such effect when IFG was stimulated.

Finally, a number of studies also measured performance changes in response to TMS or tDCS in verbal fluency tasks (see also Horvath et al., 2015; Price et al., 2015). In these tasks, participants are asked to produce as many words as possible from a given semantic category (i.e., semantic fluency) or starting with a given letter (i.e., letter fluency) within a time constraint. High fluency scores reflect unimpaired speech production on the semantic or phonological level, respectively. Neuroimaging evidence has shown that both tasks involve left frontal, temporal, and parietal regions, with dissociable activity in the MTG in the semantic and in the IFG in the letter fluency task (Birn et al., 2010). Previous studies investigating the effect of tDCS on verbal fluency have provided ambiguous results. While some studies report increased verbal fluency during or after tDCS (IFG: Cattaneo, Pisoni, & Papagno, 2011; Iyer et al., 2005; Penolazzi, Pastore, & Mondini, 2013; Pisoni, Mattavelli, et al., 2017; DLPFC: Vannorsdall et al., 2012), others did not obtain such an effect (IFG: Ehlis, Haeussinger, Gastel, Fallgatter, & Plewnia, 2016; Vannorsdall et al., 2016; DLPFC: Cerruti & Schlaug, 2009).

To date, there are still many unknowns about the influence of different stimulation parameters on the behavioural (language production) effect induced by NIBS. In order to quantify the overall effect of NIBS observed across studies and to examine individual subsets contrasting different experimental parameters, we performed a meta-analysis evaluating the behavioural performance changes during language production tasks in healthy participants. With respect to language production, rather small effect sizes of tDCS treatment for clinically relevant populations (Hartwigsen & Siebner, 2013) raise the question whether this method is a useful tool in altering language production in healthy speakers, and previous meta-analyses which investigated fewer studies are inconclusive (Horvath et al., 2015; Price et al., 2015; Westwood & Romani, 2017), as they analysed fewer studies and used diverging methods. Here, unlike these previous studies, we investigated the absolute effect sizes obtained by the application of tDCS or TMS. The direction of behavioural effects caused by NIBS (i.e., improving or disrupting performance) is difficult to predict. For instance, TMS across left temporal and inferior parietal regions has been shown to both enhance (Acheson et al., 2011; Mottaghy et al., 1999; Sparing et al., 2001; Töpper et al., 1998) and impede picture naming performance (Pobric et al., 2007, 2010; Schuhmann et al., 2012). Furthermore, the dissociation of performance improvement in response to anodal tDCS as opposed to performance decline in response to cathodal tDCS as documented for the motor cortex (Nitsche & Paulus, 2000) has been shown to be more complex for higher cognitive functions (Hill, Fitzgerald, & Hoy, 2016; Jacobson, Koslowsky, & Lavidor, 2012). For example, Fertonani et al. (2010) found (descriptive) interference in picture naming from cathodal tDCS in Experiment 1, but (descriptive) facilitation from cathodal tDCS in Experiment 2. Furthermore, a recent study reported significant facilitation from cathodal tDCS across the left pSTG in a lexical decision task (Brückner & Kammer, 2017). Given these inconsistent result patterns in both TMS and tDCS studies, we centred this meta-analysis on the question whether NIBS changes *overall* performance compared to a baseline condition, regardless of whether this change is positive or negative. Furthermore, to our knowledge, no meta-analysis has yet quantified the efficacy of TMS on inducing changes in language production in healthy speakers. Finally, by contrasting subsets of studies in regard to a number of methodological aspects (i.e., stimulation site, control condition, experimental tasks, online vs. offline stimulation, and current density of the target electrode), we intend to investigate more detailed aspects of applying NIBS in healthy speakers.

## Methods

### Study selection and analysis

To find eligible studies, we first conducted a literature search in PubMed, querying for the any combination of the search terms (“language”) AND (“tDCS”, “TMS”, “transcranial direct current stimulation”, OR “transcranial magnetic stimulation”) published up until January 2018. Additionally, the reference lists of previous reviews and meta-analyses (Hartwigsen, 2015; Horvath et al., 2015; Monti et al., 2013; Price et al., 2015) were screened to avoid overlooking suitable studies. Figure 1 provides a flowchart of the different phases of the relevant literature search.

**Figure 1.**
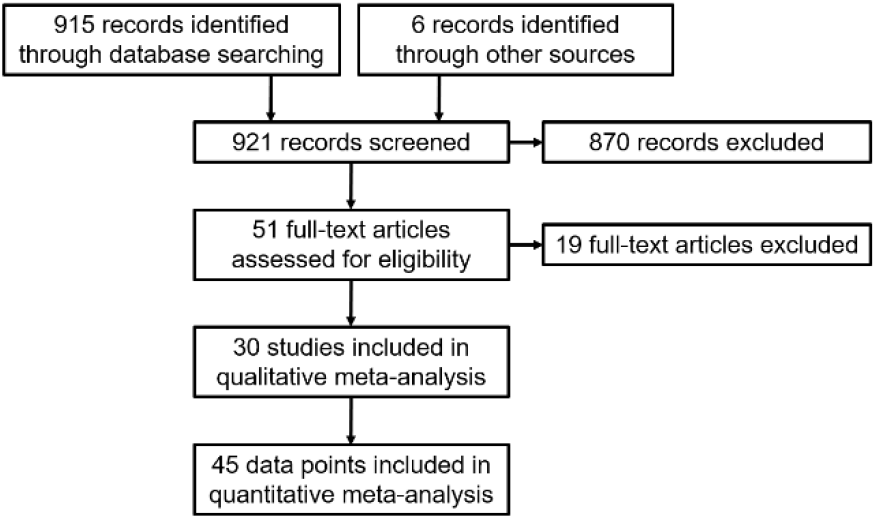
Flowchart of literature search for the meta-analysis.

Eligibility criteria were the following:

1. A single session of tDCS or TMS was applied to the left hemisphere of the cerebral cortex in right-handed participants;
2. Participants were adult healthy, young native speakers;
3. The main dependent variable was either naming latency in a picture naming task or number of words generated in a verbal fluency task;
4. The stimuli were either categories or letters (for the verbal fluency tasks), or pictures triggering single-word utterances (i.e., nouns or verbs, for picture-naming tasks). Studies using printed words as stimuli were omitted in order to avoid potential confounds with reading ability, as were studies that required the production of multi-word utterances or in which a mixture of verbal fluency and picture naming was used;
5. All relevant data were provided either in the paper or by the authors upon request, or could be extracted from figures in the publication;
6. The article was published in a peer-reviewed English-language journal;
7. The study was approved by a medical ethical committee or review board.

### Data synthesis and analysis

The literature search identified 30 eligible studies. Studies of which the full texts were screened, but which did not meet the inclusion criteria, are listed in Supplementary Table 1, along with a reason for their exclusion. For the eligible studies, the means, standard deviations, and sample sizes for all experimental and control conditions were collected (naming latencies for the picture naming tasks and number of words generated for the verbal fluency tasks). If this information was provided in graphs rather than tables, the relevant values were extracted using the software Plot Digitizer (http://plotdigitizer.sourceforge.net/). Additionally, if the reported data were not sufficient or inconsistent, the corresponding author of the paper in question was contacted and asked to provide this information. If an experiment reported several conditions (e.g., in terms of semantic category and naming cycle for semantic blocking tasks or in terms of different distractor conditions in PWI tasks), the reported values were averaged for the stimulation and the control condition in order to receive an estimate of overall effect of stimulation. All data points were coded in terms of their treatment (TMS vs. tDCS), the control condition (sham vs. no stimulation), the stimulated brain region (IFG, MTG, STG, DLPFC, IPL, or ATL), the task used (picture naming, semantic blocking, picture-word interference, or semantic fluency), the time of NIBS application (online vs. offline), and the current density of the target electrode.

For all reported comparisons (i.e., stimulation vs. control conditions) we calculated Hedges’ *d* (Rosenberg, Adams, & Gurevitch, 2000). This is an adaptation of Hedges’ *g* (Hedges & Olkin, 1985)-calculated as the difference between the mean of the experimental condition and the mean of the control condition, divided by the pooled standard deviation - which takes into account the often low sample sizes in previously published stimulation studies by multiplying the effect size with a small sample size correction. We were interested in the magnitude of the effect so we calculated the absolute effect size values. In order to avoid entering several data points from one experiment into the analysis, effect sizes originating from a single experiment were aggregated to yield a single measure per experiment. However, if several control conditions were tested which allowed for a more specific comparison of experimental variables (e.g., comparing cathodal and anodal stimulation, or different brain regions within one experiment), separate effect sizes per experiment were entered into the analysis.

We computed the cumulative effect size (i.e., the aggregated magnitude of the included studies’ effect sizes, *Ē*) and the 95% confidence intervals (CI) using a weighted average (Hedges & Olkin, 1985). All effect sizes were entered in a random effects model. As estimates of study heterogeneity, we report *Q* and *I*² values.

All effect size calculations and summary analyses were conducted using *MetaWin* (version 2.1, Rosenberg, Adams, & Gurevitch, 2000) and the *metafor* package (version 1.9-9, Viechtbauer, 2010) in *R* (version 3.3.3, R Core Team, 2017). Additional ANOVAs were run using the *ez* package (version 4.4.0, Lawrence, 2016).

## Results

In total, 45 effect sizes originating from 30 studies including 655 healthy participants were analysed (Table 1). None of the studies reported any adverse events after applying stimulation. A significant effect of NIBS was found for behavioural performance (Z = 5.214, p < .0001), indicating that applying NIBS is capable of modulating speech production processes in healthy speakers. The overall weighted mean effect size for all included studies was 0.289 (95% CI: 0.181 - 0.398). The test for heterogeneity was not significant (Q = 25.248, *p* = .990), showing that the variance between studies was not larger than is to be expected when including random sample error. Rosenberg’s fail-safe number for all studies was 274, implying that at least 274 studies publishing null effects would be required to invalidate the significant effect of NIBS on behavioural performance in language production. Figure 2 displays the effect sizes and 95% confidence intervals for all included studies.

**Table 1.**
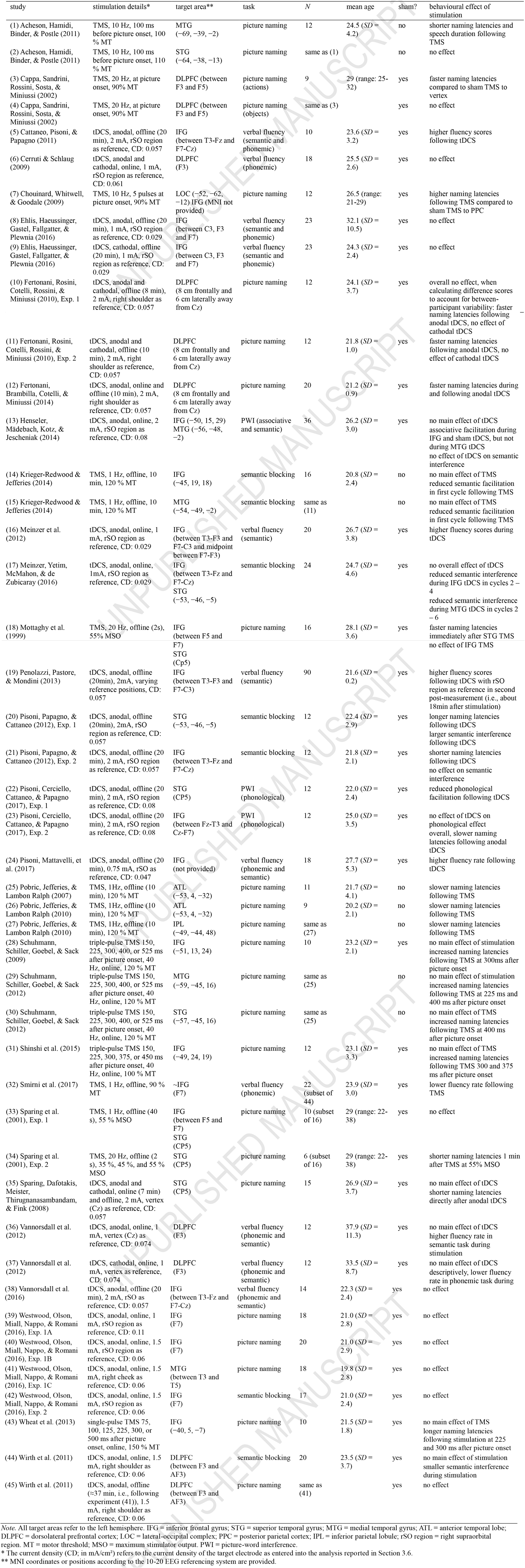
Overview of the studies included in the meta-analysis.

**Figure 2.**
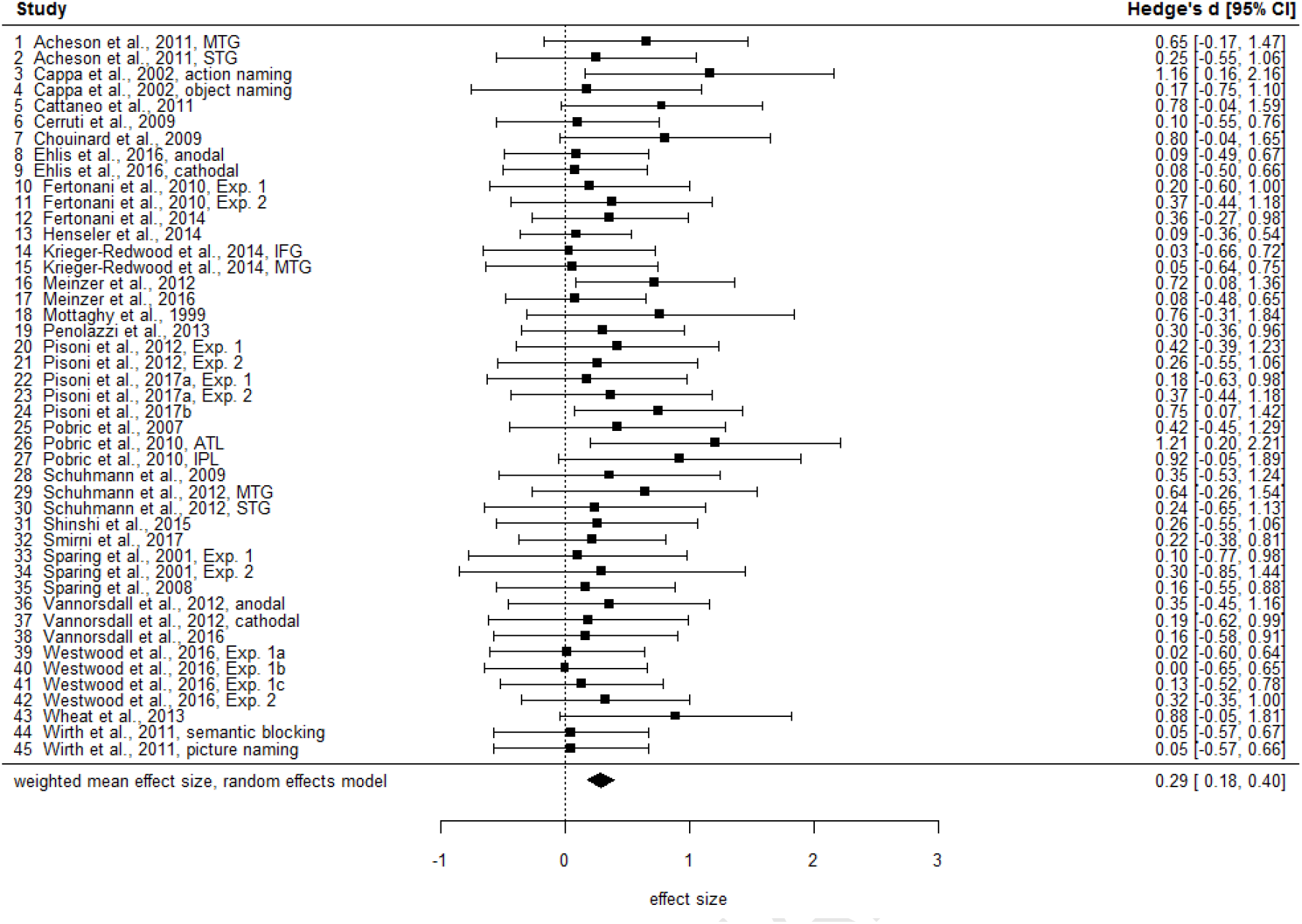
Forest plot of the effect sizes of the studies included in the meta-analysis investigating the efficacy of non-invasive brain stimulation as a tool of investigating language production in healthy participants.

Overall, our results suggest that NIBS appears to be an effective tool to modulate behaviour even in healthy participants. It should however be noted that the applied tDCS and TMS parameters used in the studies varied considerably. We therefore performed additional analyses to examine differences between stimulation type (tDCS vs. TMS), the applied control condition (sham vs. no stimulation), stimulation area (frontal vs. temporal), task (picture naming vs. verbal fluency), online vs. offline application, and current density of the target electrode. The results for these sub-analyses are summarised in Table 2.

### TMS vs. tDCS

In order to investigate the efficacy of stimulation separately for TMS (*N* = 16) and tDCS (*N* = 26), respectively, separate meta-analyses were performed for the two stimulation techniques. The outcomes revealed significant weighted mean effect sizes of 0.225 (95% CI: 0.094 - 0.356) for the tDCS studies and 0.388 (95% CI: 0.178 - 0.598) for the TMS studies (see Figure 3). Furthermore, an ANOVA comparing the effect sizes yielded a significant main effect of stimulation type (*F*(1,43) = 7.583, *p* = .009, η²G = .150), indicating that the effect sizes for the TMS studies were significantly higher than those for the tDCS studies.

**Figure 3.**
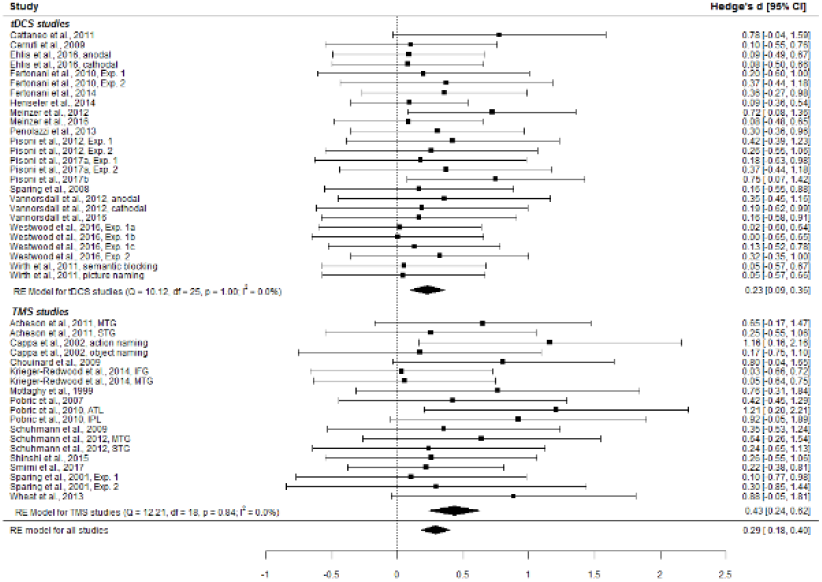
Forest plot of effect sizes, broken down by stimulation method (tDCS vs. TMS).

### Sham vs. no stimulation as a control condition

To investigate a possible difference in NIBS efficacy depending on type of control condition, we compared studies that were sham-controlled (*N* = 36) to those that were not (*N* = 9). An ANOVA yielded no significant main effect of control condition (*F*(1,43) = 2.148, *p* = .150, η²G = .048). However, separate summary analyses revealed a descriptively larger effect size for studies which were not sham-controlled (*Ē* = 0.410, 95% CI: 0.133 – 0.686) compared to those that were (*Ē* = 0.267, 95% CI: 0.149 - 0.386) (see Figure 4).

**Table 2.**
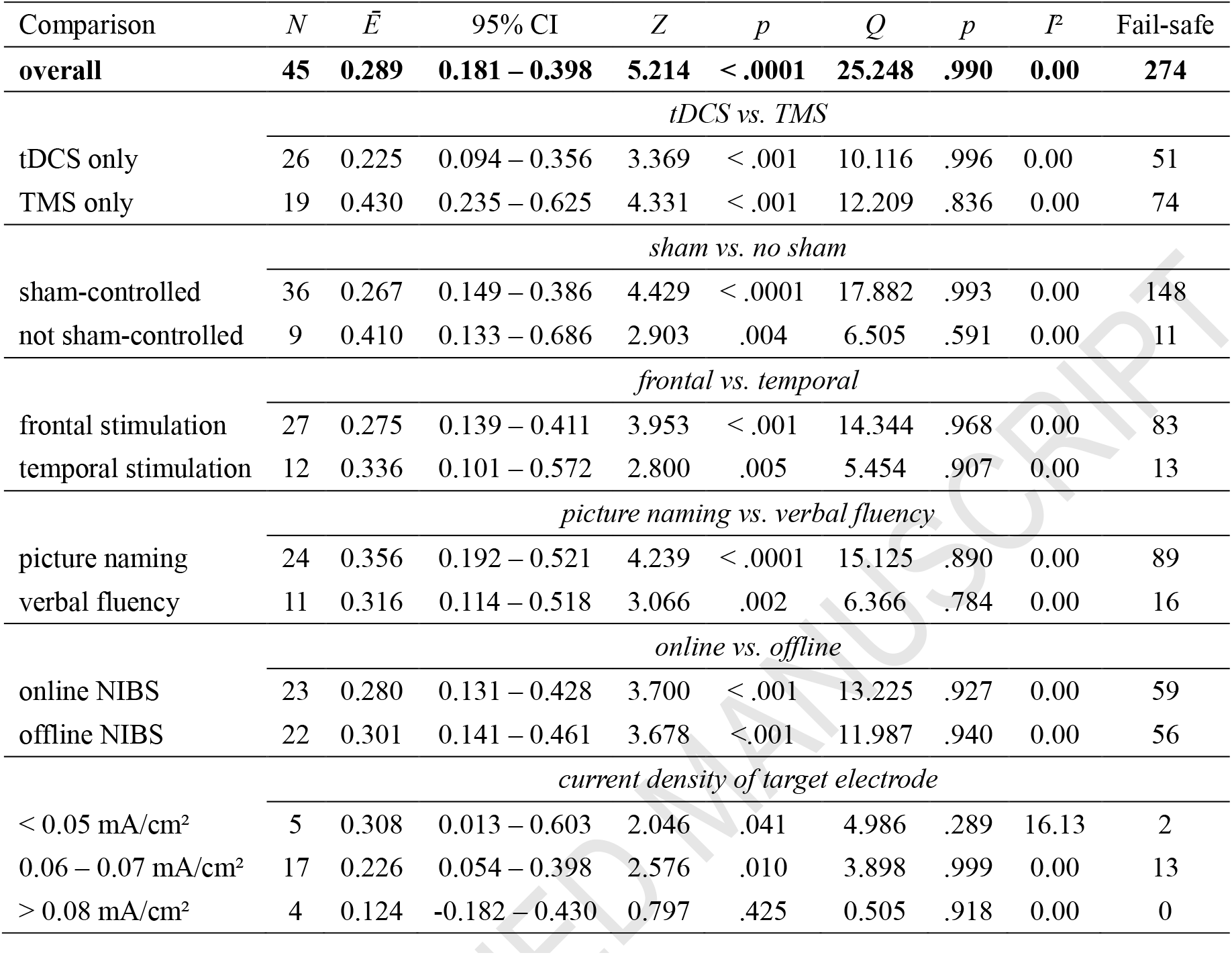
Results of meta-analysis, for all studies and specific subsets.

**Figure 4.**
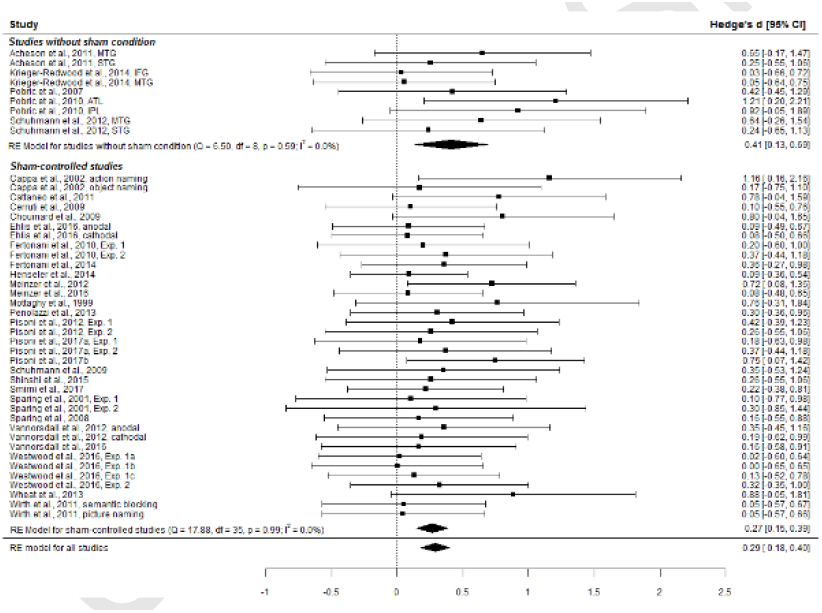
Forest plot of effect sizes, broken down by control condition (sham vs. no sham).

### Frontal vs. temporal NIBS

The majority of the studies targeted areas within the left frontotemporal language network. To investigate whether one of these regions is more susceptible to NIBS, we selected studies targeting frontal regions including the left DLPFC and the left IFG (*N* = 27), and temporal regions including the left MTG, STG, and ATL (*N* = 12). An ANOVA comparing the effect of NIBS on these two regions provided no evidence for differences in effect sizes (*F*(1,37) = 0.437, *p* = .513, η²G = .012). That is, both frontal and temporal NIBS influenced language production in healthy speakers, with no quantitative difference in the magnitude of the effect between the two target locations (for frontal regions: *Ē* = 0.275, 95% CI: 0.139 - 0.411; for temporal regions: *Ē* = 0.336, 95% CI: 0.101 - 0.572; see Figure 5).

**Figure 5.**
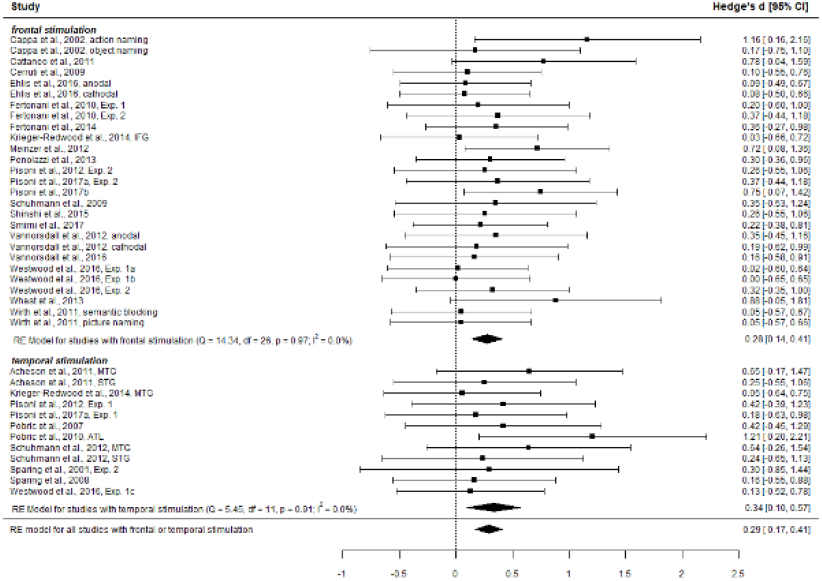
Forest plot of effect sizes, broken down by stimulation region (frontal vs. temporal).

### Picture naming vs. verbal fluency

To examine whether NIBS is more efficient for verbal fluency or picture naming tasks, we compared studies measuring verbal fluency (*N* = 11) with pure picture naming studies (*N* = 24; excluding picture-word interference and semantic blocking tasks to avoid potential confounds due to additional experimental conditions). An ANOVA provided no evidence for a difference in effect sizes between these types of tasks (*F*(1,33) = 0.597, *p* = .445, η²G = .018). Separate summary analyses yielded descriptively comparable effect sizes and confidence intervals for verbal fluency tasks (*Ē* = 0.316, 95% CI: 0.114 – 0.518) and picture naming tasks (*Ē* = 0.356, 95% CI: 0.192 – 0.521) (see Figure 6).

**Figure 6.**
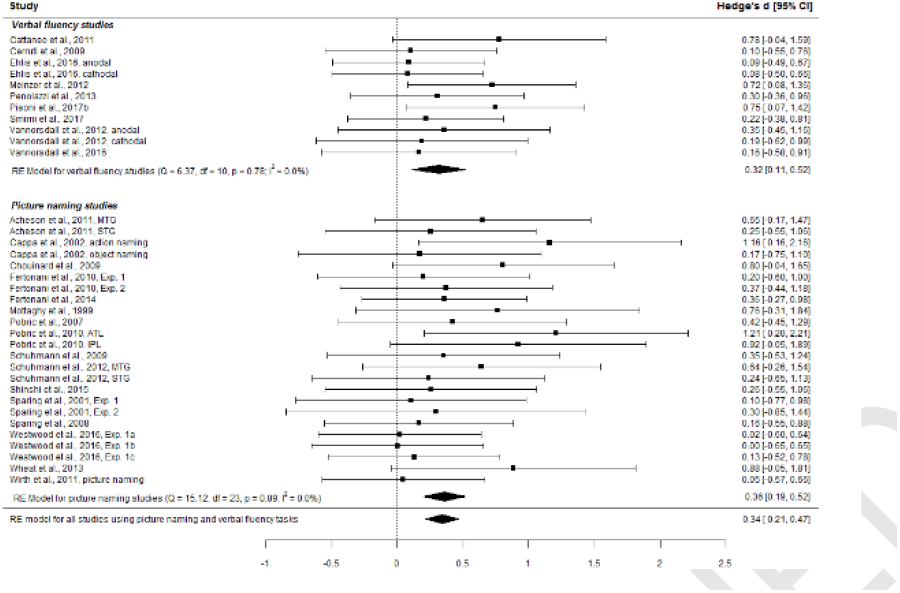
Forest plot of effect sizes, broken down by task (verbal fluency vs. picture naming).

Because frontal and temporal cortical regions are involved differentially in the tasks employed in the examined studies (see Introduction), we additionally investigated whether there are differences in effect sizes for these regions as a function of task (see Supplementary Table 2). An ANOVA including the factors region (i.e., frontal vs. temporal) and task (i.e., picture naming, semantic blocking, and picture-word interference tasks)^1^ yielded no evidence for an interaction of these two factors (*F*(2,22) = 0.175, *p* = .840, η²G = .016). However, it should be noted that sample sizes for these particular sub-analyses were very small (ranging between 2 and 11 studies). Thus, future studies are needed to allow for a more reliable estimate.

### Online vs. offline

To investigate the possible influence of applying NIBS prior to or during the execution of the experimental task, we compared studies that used an offline protocol (*N* = 22) with studies that used an online protocol (*N* = 23). An ANOVA provided no evidence for a quantitative difference between these two protocols (*F*(1,43) = 0.031, *p* = .860, η²G = .001), and effect sizes were descriptively comparable for both protocols (online: *Ē* = 0.280, 95% CI: 0.131 -0.428; offline: *Ē* = 0.301, 95% CI: 0.141 - 0.461; see Figure 7).

**Figure 7.**
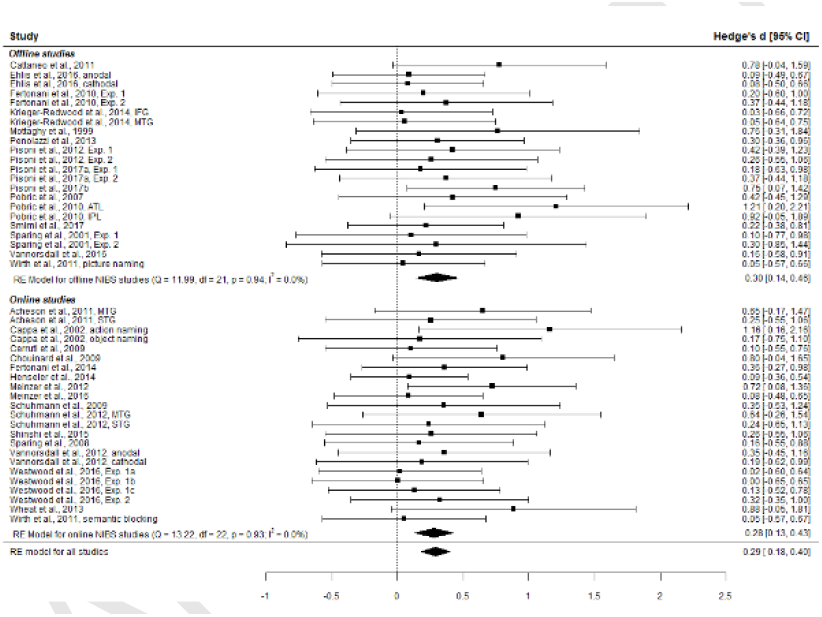
Forest plot of effect sizes, broken down by application time (online vs. offline).

### Current density of the target electrode

Finally, we investigated whether the current density of the target electrode in tDCS studies has a differential effect on performance. We treated current density as a categorical variable split in three categories (low: current density ≤ 0.05 mA/cm²; medium: current density between 0.06 and 0.07 mA/cm²; high: current density ≥ 0.08 mA/cm²). An ANOVA yielded no significant main effect of current density (*F*(2,23) = 0.747, *p* = .485, η²G = .061). Descriptively, the largest effects were observed with low (*Ē* = 0.308, 95% CI: 0.013 - 0.603) and medium current densities (*Ē* = 0.226, 95% CI: 0.054 - 0.398), whereas current densities above 0.08 mA/cm² showed no significant effect size (*Ē* = 0.124, 95% CI: -0.182 - 0.430; see Figure 8). However, it needs to be noted that most studies used current densities which we classified as “medium” (typically with a surface area between 25 and 35 cm² and a current intensity of 1.5 to 2 mA), whereas both the “low” and the “high” category are less common, thus reducing the statistical power for these groups.

**Figure 8.**
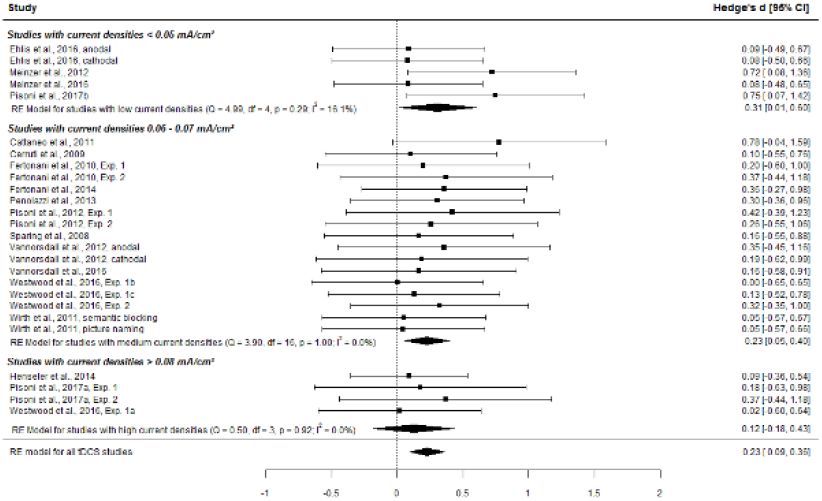
Forest plot of effect sizes, broken down by current density of active electrode (≤ 0.05 mA/cm² vs. 0.06 - 0.07 mA/cm² vs. ≥ 0.08 mA/cm²).

## Discussion

The current meta-analysis evaluated the efficacy of non-invasive brain stimulation on performance changes in language production tasks in healthy speakers. As we have reviewed in the Introduction, studies which investigated the effects of NIBS on language production performance in healthy speakers show mixed results. Importantly, the methodological approaches vary substantially between studies as well, for example, with respect to the stimulation technique, site, duration, control condition and behavioural paradigm. While there is study-specific evidence for the efficacy of NIBS in language production research, the methodological variability between studies is large. As a result, it is not clear to what extent these differences affect the behavioural outcome.

To this end, we meta-analysed the effect sizes from studies measuring picture naming latencies or verbal fluency scores in healthy participants in which either TMS or tDCS was applied to probe the causal involvement of specific cortical areas in unimpaired language production. The overall effect size for all studies combined was small, but comparable to the results found in other meta-analyses investigating the influence of NIBS on cognitive function in healthy participants (e.g., Brunoni & Vanderhasselt, 2014; Dedoncker, Brunoni, Baeken, & Vanderhasselt, 2016; Hill, Fitzgerald, & Hoy, 2016; Mancuso, Ilieva, Hamilton, & Farah, 2016; Schutter & Wischnewski, 2016). A potential reason for this relatively small effect size is that no clear-cut experimental standards exist. This introduces a large methodological variability between studies, which hampers both their comparability as well as the efficacy of the stimulation to effectively induce performance changes. For instance, for TMS studies, no valid threshold procedure (like motor-evoked potentials for the motor cortex or phosphene induction for the visual cortex) exists to reliably determine individual thresholds. Previous studies on language production used stimulation intensities between 100 and 120 % of the individual motor threshold or fixed stimulation intensities for all participants. In both cases, however, it is unclear if such a measure is the most reliable way to stimulate areas outside of the motor cortex. Inducing speech arrest may be a possible way of quantifying individual “speech thresholds”. Following Pascual-Leone, Gates, & Dhuna (1991), who had successfully induced speech arrest in epileptic patients by applying rTMS to Broca’s area, Epstein et al. (1996) contrasted the effect of stimulation frequencies between 4 and 32 Hz in a counting task. They found that applying 20 or 40 pulses over a period of five seconds (i.e., at 4 and 8 Hz, respectively) allowed for the induction of complete speech arrest without excessive muscle disturbances or pain sensations of the participants, which led the authors to conclude that this frequency was suitable for widespread application, e.g., to measure speech lateralization (see also Epstein et al., 1999). However, to the best our knowledge, none of the TMS studies that investigated language production in healthy participants has used this procedure. Similarly, for tDCS studies, individual cortical susceptibility to stimulation may differ (Parazzini, Fiocchi, Liorni, & Ravazzani, 2015), inducing different levels of excitability between participants. Also, the placement of the reference electrode, the size of both the target and reference electrode, as well as the stimulation frequencies vary substantially between studies, which hampers comparability between studies because different montages and intensities cause different electric field distributions across the cortex (Bastani & Jaberzadeh, 2013; Bastani, Jaberzadeh, Paulus, Rothwell, & Lemon, 2013; Bikson et al., 2017; Bikson, Datta, Rahman, & Scaturro, 2010; Rampersad et al., 2014; Saturnino, Antunes, & Thielscher, 2015). Further resources should be invested to explore the parameter space that allows for a reliable modulation of production performance while reducing the amount of inter- and intraindividual variability in the response to NIBS. Crucially, our results provide no evidence that applying NIBS online vs. offline, as well as the current density of the target electrode, affect weighted effect sizes.

On another note, different tasks might be differentially sensitive to performance changes induced by NIBS. We have shown that performance in both verbal fluency and pure picture naming tasks can be effectively modulated using NIBS. However, we cannot make a conclusive point with respect to the efficacy of NIBS in more specific picture naming paradigms (i.e., PWI, semantic blocking), as we have focused our analysis on the *overall* effect of NIBS as opposed to more specific experimental conditions. Westwood & Romani (2017) provide some evidence that at least tDCS may not be useful for examining semantically specific effects during language production. However, it should be noted that their analysis is based on a small number of experiments, so clearly more studies are needed before strong conclusions can be drawn.

It needs to be noted that the apparent advantage of TMS over tDCS is confounded with the physical sensations induced by either method. While participants typically cannot reliably differentiate between verum and sham tDCS, TMS arguably induces a stronger physical sensation at the stimulation site. Although some studies use so-called placebo coils or stimulate several areas (i.e., including at least one control region which is not expected to affect the outcome), many so far have only compared performance with real TMS to performance *without* the application of TMS. Evidently, in these cases, participants know when they are being stimulated and this could bias the results. We also wish to stress that for the TMS group, we pooled studies applying low-frequency (1 Hz) rTMS with studies using high-frequency (≥ 10 Hz) single- or triple-pulse TMS, which have different effects on cortical excitability. However, further subdividing the TMS studies was not meaningful given the very small sample sizes. Despite the larger effect sizes of TMS compared to tDCS, this finding should be thus treated with caution.

In conclusion, NIBS is a viable method to investigate the relations between cortical regions and language production in healthy volunteers and can contribute to the understanding of the neurobiology underlying unimpaired language production. Nevertheless, more fundamental studies are needed to explore under which conditions its efficacy can be homogenised within and between participants. Additionally, studies applying tDCS over several sessions, as is practice in clinical studies, may provide further insights into how efficacy can be improved.

## Acknowledgments

We wish to thank Barry Gordon, Beth Jefferies, Katya Krieger-Redwood, Felix Mottaghy, Marcus Meinzer, Gottfried Schlaug, Teresa Schuhmann, Rudolf Töpper, Tracy Vannorsdall, Eric Wassermann, and Greig de Zubicaray for their support in providing additional data. This work was supported by the German Research Council (KL 2933/2).

---

**Supplementary Table 1.**
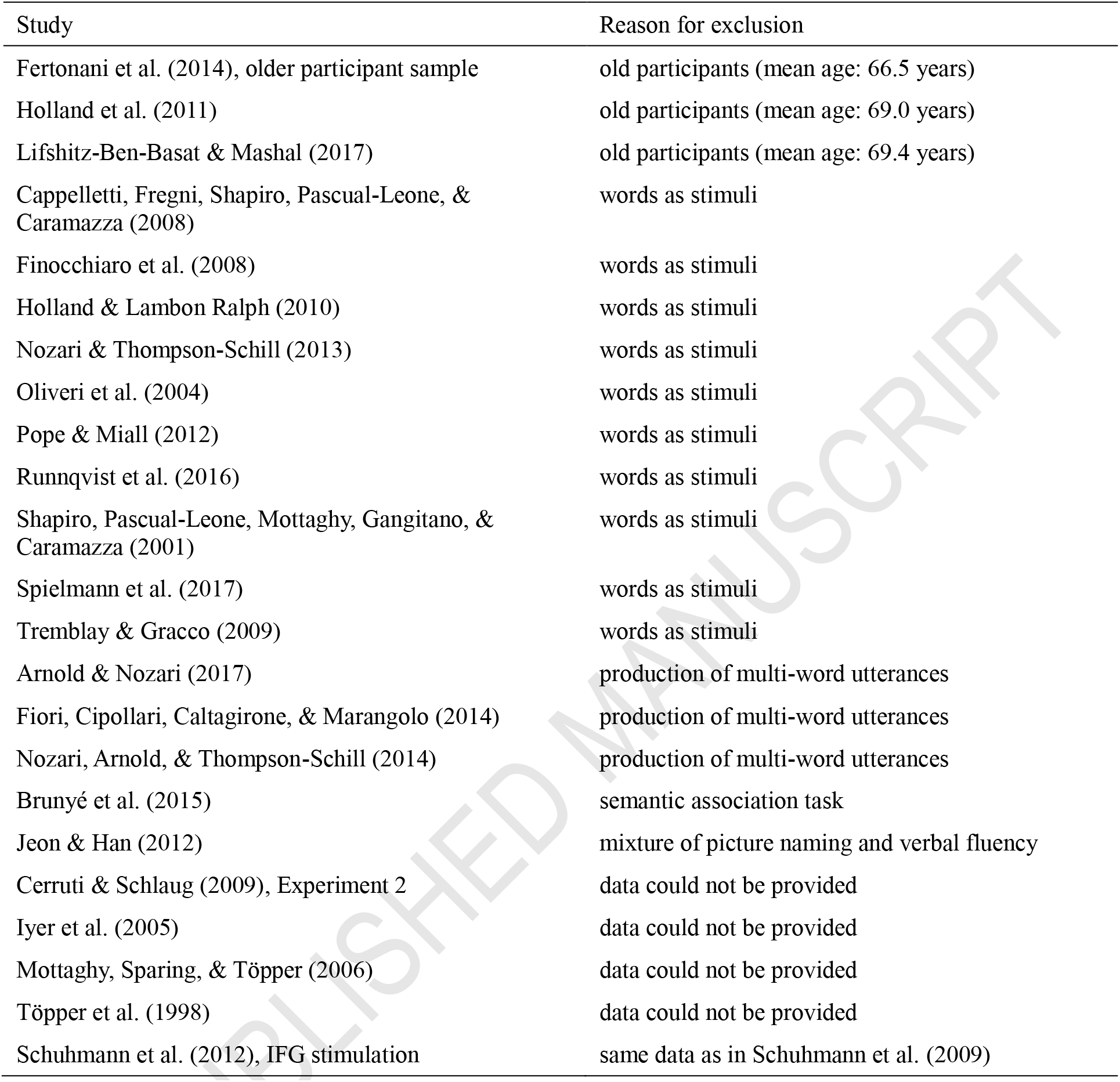
List of excluded studies and reason for exclusion.

**Supplementary Table 2.**
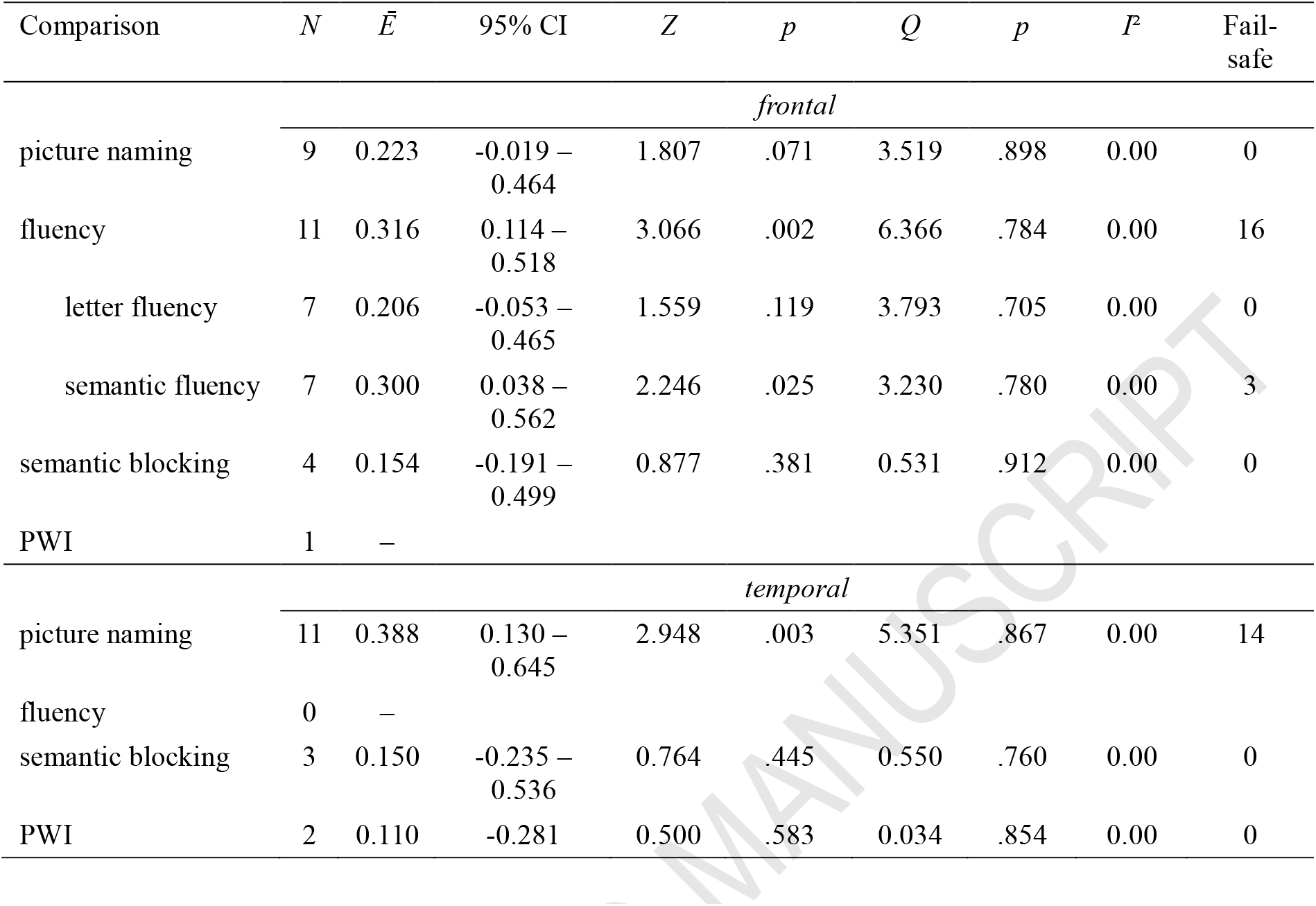
Results of sub-analysis examining the efficacy of NIBS by cortical region (frontal vs. temporal) and task.

## Footnotes

1 For this analysis, fluency tasks had to be removed because no single study has examined the effect of NIBS over temporal regions during these tasks. However, we ran additional sub-analyses for fluency tasks following frontal stimulation, both averaged across and broken down by task type (semantic vs. letter fluency). Interestingly, studies examining semantic fluency reported a stronger effect of NIBS over frontal regions, although Birn et al. (2010) showed that the left IFG is activated more during letter compared to semantic fluency tasks.

## References

Acheson, D., Hamidi, M., Binder, J., & Postle, B. (2011). A common neural substrate for language production and verbal working memory. Journal of Cognitive Neuroscience, 23(6), 1358–1367. http://doi.org/10.1162/jocn.2010.21519.

Arnold, J. E., & Nozari, N. (2017). The effects of utterance timing and stimulation of left prefrontal cortex on the production of referential expressions. Cognition, 160, 127–144. http://doi.org/10.1016/j.cognition.2016.12.008.

Bastani, A, & Jaberzadeh, S. (2013). a-tDCS Differential Modulation of Corticospinal Excitability: The Effects of Electrode Size. Brain Stimulation, 6(6), 932–937. http://doi.org/10.1016/j.brs.2013.04.005.

Bastani, A., Jaberzadeh, S., Paulus, W, Rothwell, J. C., & Lemon, R. (2013). Differential Modulation of Corticospinal Excitability by Different Current Densities of Anodal Transcranial Direct Current Stimulation. PLoS One, 8(8), e72254. http://doi.org/10.1371/journal.pone.0072254.

Belke, E., Meyer, A. S., & Damian, M. F. (2005). Refractory effects in picture naming as assessed in a semantic blocking paradigm. The Quarterly Journal of Experimental Psychology Section A, 58(4), 667–692. http://doi.org/10.1080/02724980443000142.

Bikson, M., Brunoni, A. R., Charvet, L. E., Clark, V, Cohen, L. G., Deng, Z.-D.,… Woods, A. J. (2017). Rigor and reproducibility in research with transcranial electrical stimulation: An NIMH-sponsored workshop. Brain Stimulation. http://doi.org/10.1016/j.brs.2017.12.008.

Bikson, M., Datta, A., Rahman, A., & Scaturro, J. (2010). Electrode montages for tDCS and weak transcranial electrical stimulation: role of “return” electrode’s position and size. Clinical Neurophysiology, 121(12), 1976–8. http://doi.org/10.1016/j.clinph.2010.05.020.

Birn, R. M., Kenworthy, L., Case, L., Caravella, R., Jones, T. B., Bandettini, P. A., & Martin, A. (2010). Neural systems supporting lexical search guided by letter and semantic category cues: a self-paced overt response fMRI study of verbal fluency. NeuroImage, 49(1), 1099–107. http://doi.org/10.1016/j.neuroimage.2009.07.036.

Brückner, S., & Kammer, T. (2017). Both anodal and cathodal transcranial direct current stimulation improves semantic processing. Neuroscience, 343, 269–275. http://doi.org/10.1016/J.NEUROSCIENCE.2016.12.015.

Brunoni, A. R., & Vanderhasselt, M.-A. (2014). Working memory improvement with noninvasive brain stimulation of the dorsolateral prefrontal cortex: A systematic review and meta-analysis. Brain and Cognition, 86, 1–9. http://doi.org/10.1016/j.bandc.2014.01.008.

Brunyé, T. T., Moran, J. M., Cantelon, J., Holmes, A., Eddy, M. D., Mahoney, C. R., & Taylor, H. A. (2015). Increasing breadth of semantic associations with left frontopolar direct current brain stimulation: a role for individual differences. NeuroReport, 26, 296–301. http://doi.org/10.1097/WNR.0000000000000348.

Cappa, S. F., Sandrini, M, Rossini, P. M., Sosta, K., & Miniussi, C. (2002). The role of the left frontal lobe in action naming: rTMS evidence. Neurology, 59(5), 720–723. http://doi.org/10.1212/WNL.59.5.720.

Cappelletti, M, Fregni, F., Shapiro, K. A., Pascual-Leone, A., & Caramazza, A. (2008). Processing Nouns and Verbs in the Left Frontal Cortex: A Transcranial Magnetic Stimulation Study. Journal of Cognitive Neuroscience, 20(4), 707–720. http://doi.org/10.1162/jocn.2008.20045.

Cattaneo, Z., Pisoni, A, & Papagno, C. (2011). Transcranial direct current stimulation over Broca’s region improves phonemic and semantic fluency in healthy individuals. Neuroscience, 183, 64–70. http://doi.org/10.1016/j.neuroscience.2011.03.058.

Cerruti, C, & Schlaug, G. (2009). Anodal transcranial direct current stimulation of the prefrontal cortex enhances complex verbal associative thought. Journal of Cognitive Neuroscience, 21(10), 1980–7. http://doi.org/10.1162/jocn.2008.21143.

Chouinard, P. A., Whitwell, R. L., & Goodale, M. A. (2009). The lateral-occipital and the inferior-frontal cortex play different roles during the naming of visually presented objects. Human Brain Mapping, 30(12), 3851–3864. http://doi.org/10.1002/hbm.20812.

Damian, M. F., & Martin, R. C. (1999). Semantic and phonological codes interact in single word production. Journal of Experimental Psychology: Learning, Memory and Cognition, 25, 1–18. http://doi.org/10.1037/0278-7393.25.2.345.

Damian, M. F., Vigliocco, G., & Levelt, W. J. M. (2001). Effects of semantic context in the naming of pictures and words. Cognition, 81(3), B77–B86. http://doi.org/10.1016/S0010-0277(01)00135-4.

de Zubicaray, G, Johnson, K., Howard, D., & McMahon, K. (2014). A perfusion fMRI investigation of thematic and categorical context effects in the spoken production of object names. Cortex, 54, 135–149. http://doi.org/10.1016/J.CORTEX.2014.01.018.

Dedoncker, J., Brunoni, A. R., Baeken, C, & Vanderhasselt, M.-A. (2016). A Systematic Review and Meta-Analysis of the Effects of Transcranial Direct Current Stimulation (tDCS) Over the Dorsolateral Prefrontal Cortex in Healthy and Neuropsychiatric Samples: Influence of Stimulation Parameters. Brain Stimulation, 9(4), 501–517. http://doi.org/10.1016/j.brs.2016.04.006.

Ehlis, A.-C, Haeussinger, F. B., Gastel, A., Fallgatter, A. J., & Plewnia, C. (2016). Task-dependent and polarity-specific effects of prefrontal transcranial direct current stimulation on cortical activation during word fluency. NeuroImage, 140, 134–140. http://doi.org/10.1016/j.neuroimage.2015.12.047.

Epstein, C. M., Lah, J. J., Meador, K. J., Weissman, J. D., Gaitan, L. E., & Dihenia, B. (1996). Optimum stimulus parameters for lateralized suppression of speech with magnetic brain stimulation. Neurology, 47(6), 1590–3. http://doi.org/10.1212/WNL.47.6.1590.

Epstein, C. M., Meador, K. J., Loring, D. W., Wright, R. J., Weissman, J. D., Sheppard, S.,… Davey, K. R. (1999). Localization and characterization of speech arrest during transcranial magnetic stimulation. Clinical Neurophysiology, 110(6), 1073–1079. http://doi.org/10.1016/S1388-2457(99)00047-4.

Fertonani, A., Brambilla, M., Cotelli, M., & Miniussi, C. (2014). The timing of cognitive plasticity in physiological aging: a tDCS study of naming. Frontiers in Aging Neuroscience, 6, 131. http://doi.org/10.3389/fnagi.2014.00131.

Fertonani, A., Rosini, S., Cotelli, M., Rossini, P. M., & Miniussi, C. (2010). Naming facilitation induced by transcranial direct current stimulation. Behavioural Brain Research, 208(2), 311–318. http://doi.org/10.1016/j.bbr.2009.10.030.

Finocchiaro, C, Fierro, B., Brighina, F, Giglia, G., Francolini, M., & Caramazza, A. (2008). When nominal features are marked on verbs: A transcranial magnetic stimulation study. Brain and Language, 104(2), 113–121. http://doi.org/10.1016/J.BANDL.2007.09.002.

Fiori, V, Cipollari, S., Caltagirone, C, & Marangolo, P. (2014). “If two witches would watch two watches, which witch would watch which watch?” tDCS over the left frontal region modulates tongue twister repetition in healthy subjects. Neuroscience, 256, 195–200. http://doi.org/10.1016/j.neuroscience.2013.10.048.

Hartwigsen, G. (2015). The neurophysiology of language: Insights from non-invasive brain stimulation in the healthy human brain. Brain and Language, 148, 81–94. http://doi.org/10.1016/j.bandl.2014.10.007.

Hartwigsen, G, & Siebner, H. R. (2013). Novel methods to study aphasia recovery after stroke. Frontiers in Neurology and Neuroscience, 32, 101–11. http://doi.org/10.1159/000346431.

Hedges, L. V, & Olkin, I. (1985). Statistical methods for meta-analysis. San Diego, CA: Academic Press.

Henseler, I., Mädebach, A., Kotz, S. A., & Jescheniak, J. D. (2014). Modulating Brain Mechanisms Resolving Lexico-semantic Interference during Word Production: A Transcranial Direct Current Stimulation Study. Journal of Cognitive Neuroscience, 26(7), 1403–1417. http://doi.org/10.1162/jocn_a_00572.

Hill, A. T, Fitzgerald, P. B., & Hoy, K. E. (2016). Effects of Anodal Transcranial Direct Current Stimulation on Working Memory: A Systematic Review and Meta-Analysis of Findings From Healthy and Neuropsychiatric Populations. Brain Stimulation, 9(2), 197–208. http://doi.org/10.1016/j.brs.2015.10.006.

Holland, R., & Lambon Ralph, M. A. (2010). The Anterior Temporal Lobe Semantic Hub Is a Part of the Language Neural Network: Selective Disruption of Irregular Past Tense Verbs by rTMS. Cerebral Cortex, 20(12), 2771–2775. http://doi.org/10.1093/cercor/bhq020.

Holland, R., Leff, A. P., Josephs, O., Galea, J. M., Desikan, M., Price, C. J.,… Crinion, J. (2011). Speech Facilitation by Left Inferior Frontal Cortex Stimulation. Current Biology, 21(16), 1403–1407. http://doi.org/10.1016/j.cub.2011.07.021.

Horvath, J. C, Forte, J. D., & Carter, O. (2015). Quantitative Review Finds No Evidence of Cognitive Effects in Healthy Populations From Single-session Transcranial Direct Current Stimulation (tDCS). Brain Stimulation, 8(3), 535–550. http://doi.org/10.1016/j.brs.2015.01.400.

Indefrey, P. (2011). The Spatial and Temporal Signatures of Word Production Components: A Critical Update. Frontiers in Psychology, 2, 255. http://doi.org/10.3389/fpsyg.2011.00255.

Indefrey, P., & Levelt, W. J. M. (2004). The spatial and temporal signatures of word production components. Cognition, 92(1), 101–144. http://doi.org/10.1016/j.cognition.2002.06.00 1.

Iyer, M. B., Mattu, U., Grafman, J., Lomarev, M., Sato, S., & Wassermann, E. M. (2005). Safety and cognitive effect of frontal DC brain polarization in healthy individuals. Neurology, 64(5), 872–5. http://doi.org/10.1212/01.WNL.0000152986.07469.E9.

Jacobson, L., Koslowsky, M., & Lavidor, M. (2012). tDCS polarity effects in motor and cognitive domains: a meta-analytical review. Experimental Brain Research, 216(1), 1–10. http://doi.org/10.1007/s00221-011-2891-9.

Jeon, S. Y, & Han, S. J. (2012). Improvement of the Working Memory and Naming by Transcranial Direct Current Stimulation. Ann Rehabil Med, 36(5), 585–595. http://doi.org/10.5535/arm.2012.36.5.585.

Krieger-Redwood, K., & Jefferies, E. (2014). TMS interferes with lexical-semantic retrieval in left inferior frontal gyrus and posterior middle temporal gyrus: Evidence from cyclical picture naming. Neuropsychologia, 64, 24–32. http://doi.org/10.1016/j.neuropsychologia.2014.09.014.

Kroll, J. F, & Stewart, E. (1994). Category Interference in Translation and Picture Naming: Evidence for Asymmetric Connections Between Bilingual Memory Representations. Journal of Memory and Language, 33(2), 149–174. http://doi.org/10.1006/jmla.1994.1008.

Lawrence, M. A. (2016). ez: Easy Analysis and Visualization of Factorial Experiments. R package version 4.4-0. Retrieved from https://cranr-project.org/package=ez.

Lifshitz-Ben-Basat, A, & Mashal, N. (2017). Improving Naming Abilities Among Healthy Young-Old Adults Using Transcranial Direct Current Stimulation. Journal of Psycholinguistic Research, 1–12. http://doi.org/10.1007/s10936-017-9516-9.

Mancuso, L. E., Ilieva, I. P., Hamilton, R. H., & Farah, M. J. (2016). Does Transcranial Direct Current Stimulation Improve Healthy Working Memory?: A Meta-analytic Review. Journal of Cognitive Neuroscience, 28(8), 1063–1089. http://doi.org/10.1162/jocn_a_00956.

Meinzer, M., Antonenko, D., Lindenberg, R., Hetzer, S., Ulm, L., Avirame, K.,… Flöel, A. (2012). Electrical Brain Stimulation Improves Cognitive Performance by Modulating Functional Connectivity and Task-Specific Activation. The Journal of Neuroscience, 32(5), 1859–1866. http://doi.org/10.1523/JNEUROSCI.4812-11.2012.

Meinzer, M., Yetim, Ö., McMahon, K., & de Zubicaray, G. (2016). Brain mechanisms of semantic interference in spoken word production: An anodal transcranial Direct Current Stimulation (atDCS) study. Brain and Language, 157, 72–80. http://doi.org/10.1016/j.bandl.2016.04.003.

Monti, A., Ferrucci, R., Fumagalli, M., Mameli, F, Cogiamanian, F, Ardolino, G., & Priori, A. (2013). Transcranial direct current stimulation (tDCS) and language. Journal of Neurology, Neurosurgery & Psychiatry, 84(8), 832–842. http://doi.org/10.1136/jnnp-2012-302825.

Mottaghy, F M., Hungs, M., Brügmann, M., Sparing, R, Boroojerdi, B., Foltys, H.,… Töpper, R. (1999). Facilitation of picture naming after repetitive transcranial magnetic stimulation. Neurology, 53(8), 1806–12. http://doi.org/10.1212/WNL.53.8.1806.

Mottaghy, F M., Sparing, R, & Töpper, R. (2006). Enhancing Picture Naming with Transcranial Magnetic Stimulation. Behavioural Neurology, 17(3-4), 177–186. http://doi.org/10.1155/2006/768413.

Nitsche, M. A., Cohen, L. G., Wassermann, E. M., Priori, A., Lang, N., Antal, A.,… Pascual-Leone, A. (2008). Transcranial direct current stimulation: State of the art 2008. Brain Stimulation, 1(3), 206–223. http://doi.org/10.1016/j.brs.2008.06.004.

Nitsche, M. A., & Paulus, W. (2000). Excitability changes induced in the human motor cortex by weak transcranial direct current stimulation. The Journal of Physiology, 527(3), 633–639. http://doi.org/10.1111/j.1469-7793.2000.t01-1-00633.x.

Nozari, N., Arnold, J. E., & Thompson-Schill, S. L. (2014). The effects of anodal stimulation of the left prefrontal cortex on sentence production. Brain Stimulation, 7(6), 784–92. http://doi.org/10.1016/j.brs.2014.07.035.

Nozari, N., & Thompson-Schill, S. L. (2013). More attention when speaking: Does it help or does it hurt? Neuropsychologia, 51(13), 2770–2780. http://doi.org/10.1016/j.neuropsychologia.2013.08.019.

Oliveri, M., Finocchiaro, C, Shapiro, K., Gangitano, M., Caramazza, A., & Pascual-Leone, A. (2004). All Talk and No Action: A Transcranial Magnetic Stimulation Study of Motor Cortex Activation During Action Word Production. Journal of Cognitive Neuroscience, 16(3), 374–381. http://doi.org/10.1162/089892904322926719.

Parazzini, M., Fiocchi, S., Liorni, I., & Ravazzani, P. (2015). Effect of the Interindividual Variability on Computational Modeling of Transcranial Direct Current Stimulation. Computational Intelligence and C Neuroscience, 2015, 963293. http://doi.org/10.1155/2015/963293.

Pascual-Leone, A., Gates, J. R., & Dhuna, A. (1991). Induction of speech arrest and counting errors with rapid-rate transcranial magnetic stimulation. Neurology, 41(5), 697–702. http://doi.org/10.1212/WNL.41.5.697.

Penolazzi, B., Pastore, M., & Mondini, S. (2013). Electrode montage dependent effects of transcranial direct current stimulation on semantic fluency. Behavioural Brain Research, 248, 129–135. http://doi.org/10.1016/j.bbr.2013.04.007.

Pisoni, A., Cerciello, M., Cattaneo, Z., & Papagno, C. (2017). Phonological facilitation in picture naming: When and where? A tDCS study. Neuroscience, 352, 106–121. http://doi.org/10.1016/j.neuroscience.2017.03043.

Pisoni, A., Mattavelli, G, Papagno, C, Rosanova, M., Casali, A. G., & Romero Lauro, L. J. (2017). Cognitive Enhancement Induced by Anodal tDCS Drives Circuit-Specific Cortical Plasticity. Cerebral Cortex. http://doi.org/10.1093/cercor/bhx021.

Pisoni, A., Papagno, C, & Cattaneo, Z. (2012). Neural correlates of the semantic interference effect: New evidence from transcranial direct current stimulation. Neuroscience, 223, 56–67. http://doi.org/10.1016/j.neuroscience.2012.07046.

Pobric, G, Jefferies, E., & Lambon Ralph, M. A. (2007). Anterior temporal lobes mediate semantic representation: mimicking semantic dementia by using rTMS in normal participants. Proceedings of the National Academy of Sciences of the United States of America, 104(50), 20137–41. http://doi.org/10.1073/pnas.0707383104.

Pobric, G, Jefferies, E., & Lambon Ralph, M. A. (2010). Category-Specific versus Category-General Semantic Impairment Induced by Transcranial Magnetic Stimulation. Current Biology, 20(10), 964–968. http://doi.org/10.1016/j.cub.2010.03.070.

Pope, P. A., & Miall, R. C. (2012). Task-specific facilitation of cognition by cathodal transcranial direct current stimulation of the cerebellum. Brain Stimulation, 5, 84–94. http://doi.org/10.1016/j.brs.2012.03.006.

Price, A. R., McAdams, H., Grossman, M, & Hamilton, R. H. (2015). A Meta-analysis of Transcranial Direct Current Stimulation Studies Examining the Reliability of Effects on Language Measures. Brain Stimulation, 8(6), 1093–1100. http://doi.org/10.1016/j.brs.2015.06.013.

Rampersad, S. M., Janssen, A. M., Lucka, F., Aydin, U., Lanfer, B., Lew, S.,… Oostendorp, T. F. (2014). Simulating Transcranial Direct Current Stimulation With a Detailed Anisotropic Human Head Model. IEEE Transactions on Neural Systems and Rehabilitation Engineering, 22(3), 441–452. http://doi.org/10.1109/TNSRE.2014.2308997.

Rosenberg, M. S., Adams, D. C., & Gurevitch, J. (2000). MetaWin. Statistical Software for Meta-Analysis. Sunderland, MA: Sinauer Associates.

Runnqvist, E., Bonnard, M., Gauvin, H. S., Attarian, S., Trébuchon, A., Hartsuiker, R. J., & Alario, F.-X. (2016). Internal modeling of upcoming speech: A causal role of the right posterior cerebellum in non-motor aspects of language production. Cortex, 81, 203–214. http://doi.org/10.1016/j.cortex.2016.05.008.

Saturnino, G B., Antunes, A., & Thielscher, A. (2015). On the importance of electrode parameters for shaping electric field patterns generated by tDCS. NeuroImage, 120, 25–35. http://doi.org/10.1016/j.neuroimage.2015.06.067.

Schriefers, H, Meyer, A. S., & Levelt, W. J. M. (1990). Exploring the time course of lexical access in language production: Picture-word interference studies. Journal of Memory and Language, 29(1), 86–102. http://doi.org/10.1016/0749-596X(90)90011-N.

Schuhmann, T, Schiller, N. O., Goebel, R, & Sack, A. T (2009). The temporal characteristics of functional activation in Broca’s area during overt picture naming. Cortex, 45(9), 1111–1116. http://doi.org/10.1016/j.cortex.2008.10.013.

Schuhmann, T, Schiller, N. O., Goebel, R, & Sack, A. T (2012). Speaking of which: dissecting the neurocognitive network of language production in picture naming. Cerebral Cortex, 22(3), 701–709. http://doi.org/10.1093/cercor/bhr155.

Schutter, D. J. L. G, & Wischnewski, M. (2016). A meta-analytic study of exogenous oscillatory electric potentials in neuroenhancement. Neuropsychologia, 86, 110–118. http://doi.org/10.1016/j.neuropsychologia.2016.04.011.

Shapiro, K. A., Pascual-Leone, A., Mottaghy, F. M., Gangitano, M., & Caramazza, A. (2001). Grammatical Distinctions in the Left Frontal Cortex. Journal of Cognitive Neuroscience, 13(6), 713–720. http://doi.org/10.1162/08989290152541386.

Shinshi, M., Yanagisawa, T, Hirata, M., Goto, T, Sugata, H, Araki, T,… Yorifuji, S. (2015). Temporospatial identification of language-related cortical function by a combination of transcranial magnetic stimulation and magnetoencephalography. Brain and Behavior, 5(3), e00317. http://doi.org/10.1002/brb3.317.

Smirni, D., Turriziani, P., Mangano, G. R., Bracco, M., Oliveri, M., & Cipolotti, L. (2017). Modulating phonemic fluency performance in healthy subjects with transcranial magnetic stimulation over the left or right lateral frontal cortex. Neuropsychologia, 102, 109–115. http://doi.org/10.1016/j.neuropsychologia.2017.06.006.

Sparing, R, Dafotakis, M., Meister, I. G, Thirugnanasambandam, N., & Fink, G. R. (2008). Enhancing language performance with non-invasive brain stimulation—A transcranial direct current stimulation study in healthy humans. Neuropsychologia, 46(1), 261–268. http://doi.org/10.1016/j.neuropsychologia.2007.07.009.

Sparing, R, Mottaghy, F. M., Hungs, M., Bruegmann, M., Foltys, H, Huber, W., & Töpper, R. (2001). Repetitive transcranial magnetic stimulation effects on language function depend on the stimulation parameters. Journal of Clinical Neurophysiology, 18(4), 326–330. http://doi.org/10.1097/00004691-200107000-00004.

Spielmann, K., van der Vliet, R, van de Sandt-Koenderman, W. M. E., Frens, M. A., Ribbers, G. M., Selles, R. W.,… Holland, P. (2017). Cerebellar Cathodal Transcranial Direct Stimulation and Performance on a Verb Generation Task: A Replication Study. Neural Plasticity, 2017, 1–12. http://doi.org/10.1155/2017/1254615.

Stagg, C. J., & Nitsche, M. A. (2011). Physiological Basis of Transcranial Direct Current Stimulation. The Neuroscientist, 17(1), 37–53. http://doi.org/10.1177/1073858410386614.

Töpper, R., Mottaghy, F. M., Brügmann, M., Noth, J., & Huber, W. (1998). Facilitation of picture naming by focal transcranial magnetic stimulation of Wernicke’s area. Experimental Brain Research, 121(4), 371–378. http://doi.org/10.1007/s002210050471.

Tremblay, P., & Gracco, V. L. (2009). Contribution of the pre-SMA to the production of words and non-speech oral motor gestures, as revealed by repetitive transcranial magnetic stimulation (rTMS). Brain Research, 1268, 112–124.

Vannorsdall, T. D., Schretlen, D. J., Andrejczuk, M, Ledoux, K., Bosley, L. V., Weaver, J. R.,… Gordon, B. (2012). Altering automatic verbal processes with transcranial direct current stimulation. Frontiers in Psychiatry, 3, 73. http://doi.org/10.3389/fpsyt.2012.00073.

Vannorsdall, T. D., van, Steenburgh J. J., Schretlen, D. J., Jayatillake, R., Skolasky, R. L., & Gordon, B. (2016). Reproducibility of tDCS Results in a Randomized Trial. Cognitive And Behavioral Neurology, 29(1), 11–17. http://doi.org/10.1097/WNN.0000000000000086.

Viechtbauer, W. (2010). Conducting meta-analyses in R with the metafor package. Journal of Statistical Software, 36(3), 1–48.

Westwood, S. J., Olson, A, Miall, R. C., Nappo, R, & Romani, C. (2017). Limits to tDCS effects in language: Failures to modulate word production in healthy participants with frontal or temporal tDCS. Cortex, 86, 64–82. http://doi.org/10.1016/j.cortex.2016.10.016.

Westwood, S. J., & Romani, C. (2017). Transcranial direct current stimulation (tDCS) modulation of picture naming and word reading: A metaanalysis of single session tDCS applied to healthy participants. Neuropsychologia. http://doi.org/10.1016/j.neuropsychologia.201 7.07.031.

Wheat, K. L., Cornelissen, P. L., Sack, A. T., Schuhmann, T, Goebel, R, & Blomert, L. (2013). Charting the functional relevance of Broca’s area for visual word recognition and picture naming in Dutch using fMRI-guided TMS. Brain and Language, 125(2), 223–230. http://doi.org/10.1016/j.bandl.2012.04.016.

Wirth, M, Rahman, R. A., Kuenecke, J., Koenig, T, Horn, H., Sommer, W., & Dierks, T (2011). Effects of transcranial direct current stimulation (tDCS) on behaviour and electrophysiology of language production. Neuropsychologia, 49(14), 3989–3998. http://doi.org/10.1016/j.neuropsychologia.2011.10.015.

